# The detection of covariation of mRNA levels of large sets of genes across multiple human populations

**DOI:** 10.1101/082842

**Authors:** Yu Quan, Chao Xie, Rohan B. H. Williams, Peter F. R Little

**Author notes:** Peter F. R. Little, Life Sciences Institute (LSI), National University of Singapore, Centre for Life Sciences, #05-02, 28 Medical Drive, Singapore 117456.

## Abstract

In this study, we analyse RNA-Seq data from panels of human lymphoblastoid cell lines (LCLs) to identify covariation in the mRNA levels of large numbers of genes. Such large scale covariation may have biological origin or be due to technical variation in analysis (generally referred to as batch effects). We show that batch effects cannot explain this covariation by demonstrating reproducibility across different human populations and across different methods of analysis. This view is also supported by enrichment of single and combinations of transcription factors (TFs) binding to cognate promoter regions, enrichment of genes shown to be sensitive to the knockdown of individual TFs, enrichment of functional pathways, and finally enrichment of protein-protein interactions in proteins encoded by groups of covarying genes. The properties of the groups of covarying genes are therefore most readily explained by the influence of cumulative variations in the effectors of gene expression that act in *trans* on cognate genes. We suggest that covariation has functional outcomes by showing that covariation of 83 genes involved in the spliceosome pathway accounts for 8–16% of the variation in the alternative splicing patterns of genes expressed in human LCLs.

## Introduction

Genetic variation that influences gene expression in humans is thought to be a major mechanism contributing to human phenotypes.^1^ The variation in mRNA levels in normal human cells is very substantial. The mRNA levels of around 14,000 genes expressed and measured by mRNA-Seq in lymphoblastoid cell lines (LCLs) derived from 88 Yoruban individuals (data from Lappalainen et al.^2^) has a median of 4.3 fold change by comparing the minimum to maximum expression levels. Presently, the overall inter-individual genetically derived variation in the expression level of a gene is best explained by a mix of *cis*- and *trans*-acting eQTLs (expression quantitative trait loci) defined as associated genetic variations local (*cis*) or distant (*trans*) to the affected gene based on the observation that local eQTLs usually act in *cis* and distant eQTLs usually act in *trans*.^3^ Huan et al.^4^ reported that *cis*-acting eQTLs account for 33–53% of mRNA variability of a single gene whereas individual *trans*-eQTLs explain only 2–7%. This information derives from detecting the influence of a single eQTL on a single gene, be it in *cis* or in *trans*, with *trans-*eQTLs being defined simply by virtue of having a remote location from the cognate gene. However, it is also clear that the proteins and RNA elements involved in *trans*-acting controls on gene expression invariably have multiple gene or mRNA targets, and so a common feature of variation in *trans* influences should be that they will cause simultaneous changes to the mRNA levels of multiple genes.^5^ The impact of variation in these pleiotropic regulators can be substantial; Lovén et al.^6^ showed that engineered overexpression of the *MYC* gene results in increases in expression of 90% of all genes expressed in the cells and Cusanovich et al.^7^ used individual RNAi knockdowns of 59 transcription factors (TFs) to show that between 39 and 3,892 genes were differentially expressed as a consequence of knockdown of individual TFs. The knockdown of each TF mRNA level ranged 52–92% of normal in the target LCL, and analysis of mRNA encoding TFs measured in LCLs of multiple normal human beings reveals very similar levels of variation (Figure S1). This observation suggests that variation in TF mRNA levels in normal humans might result from in *trans* influences upon multiple genes. Moreover, genetic variations on coding sequences of TFs can affect their DNA-binding activity.^8^ Such simultaneous variation could be of substantial importance to phenotypic variation in humans, simply because many *trans* controls, particularly TFs, have been implicated in control of genes organised within pathways of function,^7,9–12^ suggesting that *trans* variation might have pathway specific influences.

Detecting *trans* genetic influences presents two substantial challenges to conventional eQTL strategies: firstly, very large sample sizes are required to overcome the multiple testing problem implicit in analysing each mRNA trait against all genomic SNPs, and secondly the influence of a single *trans*-eQTL upon a target gene is weak.^4^ An alternative to the gene/eQTL centric approach is to infer the outcome of *trans* influence by detecting the correlated variation of the mRNA levels of groups of target genes in multiple individuals.^10,13^ Covariation of mRNA levels in such an analysis will necessarily be due to cumulative effects of trans-acting variations, be they caused by TFs or any other diffusible regulators, and the detection of these overall effects does not lead to the identification of the underlying *trans* eQTLs but rather identifies the cumulative influence of all *trans* influences on mRNA levels within the individuals under study. Importantly, detecting covariation of mRNA levels in a set of genes as an analytic approach is statistically tractable in extant sample sets because it is not confounded by the *trans*-eQTL multiple testing problem, nor is it necessarily limited to identifying the small effect associated with single *trans*-eQTLs.

However, the major difficulty in detecting covariation due to *trans* influences is to distinguish this from the covariation in mRNA levels that is induced by a wealth of technical artefacts, collectively referred to as batch effects, which are inherent to the analytic methods used in mRNA level estimation. Such effects include systematic differences in mRNA isolation and yield, purity, cDNA library or target mRNA construction and ascertainment of yield by both mRNA-Seq and by microarrays.^14^ Many methods have been developed^15^ to control for such batch effects but in most cases they are explicitly designed to detect and eliminate large-scale covariation in mRNA levels since these methods are intended to provide accurate mRNA measure on a gene by gene basis. If *trans* influences are indeed present on human gene expression then the likely outcome would be covariation of mRNA levels of multiple genes, that based upon the data of Cusanovich et al.,^7^ could number in the thousands. Goldinger et al.^16^ used eQTL methodology to identify *trans* effects in human mRNA samples analysed by microarrays and were able to show that principal component analysis (PCA), a statistical procedure commonly used to control batch effects, could also remove *trans* genetic influences. In this paper, we report an analysis based upon detection of large scale covariation of mRNA level using PCA to detect simultaneous variation of large sets of genes across multiple normal individuals. Following the previous analysis of Cowley et al.,^13^ we will refer to these sets of genes as *correlating group of genes*, abbreviated to CGG.

We show that the genes within CGGs share multiple biological properties and that this makes it unlikely that covariation is simply a product of batch effects. Our observations lead us to suggest that covariation of mRNA levels in sets of human genes is common and could contribute directly to human phenotypic variability at both the individual and population level. Further, we show the covariation of a set of 83 genes that are involved in mRNA splicing has a significant influence upon splicing patterns of human genes in LCLs.

## Materials and Methods

### Datasets of mRNA level in human LCLs

1. Lappalainen et al:^2^ mRNA-Seq data from LCLs using the Illumina HiSeq2000 platform. Reads per kilobase per million mapped reads (RPKM) values of genes expressed in LCLs from Caucasian (CEU), Yoruba (YRI), Finns (FIN), British (GBR) and Toscani (TSI) individuals were obtained from ArrayExpress: E-GEUV-1.^17^ We selected expressed genes defined as genes with non-zero RPKM values across all LCL samples yielding 15,016 genes from CEU, 14,918 genes from YRI and 15,231 genes from FIN+GBR+TSI. Where available, RPKM values of duplicate samples from the same LCL were averaged for each gene. We inspected log10 transformed RPKM distributions for all samples and excluded 1 YRI, 2 FIN and 1 GBR samples with outlier distribution, resulting in RPKM values of 88 individuals from YRI, 91 individuals from CEU and 279 individuals from FIN+GBR+TSI. 88 individuals were then randomly selected from CEU and FIN+GBR+TSI dataset to make 3 datasets with the same sample size. To correct for confounding effect of gender on mRNA level measurements, linear regression of each mRNA level profile on gender was then applied using R function *lm*.
2. Pickrell et al:^18^ mRNA-Seq data from LCLs from 69 Yoruba individuals sequenced at Yale (YRI Yale) and Argonne (YRI Argonne) sequencing center using the Illumina GAM platform (GEO: GSE19480), and Montgomery et al:^19^ mRNA-Seq data of LCLs from 60 Caucasians (CEU) using the Illumina GAM platform (ArrayExpress: E-MTAB-197). To quantify mRNA level of genes, for both RNA-Seq data from Pickrell et al.^18^ and Montgomery et al.^19^ reads were mapped to the human reference genome GRCh37 using BWA^20^ and Samtools,^21^ and then counted using Rsamtools^22^ *and* GenomicRanges^23^ based on human gene annotations from Ensembl Genes 69.^24^ RPKM values were then calculated for each gene. We detected expressed genes including 12,171 genes from YRI Yale data, 12,385 genes from YRI Argonne data, and 9,418 genes from CEU data. RPKM values of duplicate samples from the same LCL were averaged for each gene. We inspected log1O transformed RPKM distributions for all samples and excluded 1 YRI Yale, 1 YRI Argonne and 6 CEU samples with outlier distribution. RPKM values with the remaining 68 individuals from YRI Yale data, the same 68 individuals from YRI Argonne data, and 54 individuals from CEU data were included in the following analysis. Linear regression of each mRNA level profile for YRI Yale and Argonne data was carried out against sample RNA concentration and gender, and for CEU data against gender, in both cases using R function *lm*.
3. Stranger et al:^25^ gene expression microarray data from LCLs from 109 Caucasian individuals using Illumina’s Human-6 Expression BeadChip version 2. Normalized mRNA level values were obtained from ArrayExpress: E-MTAB-198. We detected 15,211 expressed genes from the microarray mRNA level dataset. Linear regression of each mRNA level profile on gender was performed using R function *lm*.

### Principal component analysis

To calculate mRNA level matrices *X* with sequentially removed principal components (PCs), we scaled each log10 transformed mRNA level profile into mean 0 and standard deviation 1, and then performed PCA on mRNA level data for each dataset using R function *svd*. After obtaining *X*=*UDV^T^*, we performed filtering of the first *k* PCs by setting the first *k* diagonal elements of the D matrix to zero, denoted as *D_k_*, and then calculated *X_k_* using *X_k_*=*UD_k_V^T^*, so that the variance explained by the first *k* PCs was removed from mRNA level matrix *X*.

### mRNA correlation analysis

We calculated correlations of all gene pairs from all mRNA level matrices using absolute Spearman’s correlation coefficient |ρ| computed via the R function *cor*. To generate the null distribution of correlation, 1000 permutations were applied to each mRNA level matrix by randomly permuting the individual labels of each gene expression profile across individuals. Gene correlations with the permutation p-value < 0.01 were considered as significant (Figure S2 and Table S1). Gene correlation networks were then built by connecting gene pairs with correlation above the threshold by edges using function *graph.edgelist* from igraph.^26^

### Simulation of batch effects

To investigate the influence of batch effects on the detection of co-varying genes, we simulate artificial batch effects by increasing gene expression level of random sets of genes in random 44 out of 88 RNA-Seq samples. All RPKM values were first log10 transformed and normalized to have 0 mean and unit 1 standard deviation (SD), and then an offset (0.5,1.0,1.5 or 2.0 SDs) were added to expression level of random genes (200, 500, 1000 or 2000 genes) in random 44 samples. We found that adding offset of 0.5, 1.0, 1.5 or 2.0 SDs correspond to 1.15, 1.33, 1.54 and 1.78 fold change of RPKM values, respectively. After artificially increasing mRNA levels of certain genes, we applied the same procedure as described in the main text to identify the combined CGG. The same simulation process was applied on two gene expression datasets: one is RNA-Seq data of LCLs from Yoruba individuals,^2^ the other one is the same dataset but with sample labels randomly permutated for each gene expression profile. The simulation process was repeated 100 times, and then the median value of summary statistics was recorded.

### Replication analyses

We used Fisher’s exact test to test whether there is significant overlap of genes between CGGs identified from different data sets using R function *fisher.test*, Bonferroni corrected for the number of comparisons that were performed.

### PEER and GC-content bias correction

PEER was used in processing mRNA level data with 22 unobserved factors (25% of sample size) as recommended in Stegle et al.^27^ To investigate the effect of GC-content bias on the covariation of mRNA level we adapted the GC-content bias correction procedure as described in Pickrell et al.^18^ All expressed genes were first grouped into 200 bins of equal size based on their gene-level GC content. Then, the log2 relative enrichment of RPKM values was calculated for each gene bin from each sample. We then fitted a smoothing spline for the relative enrichment of each gene bin against its mean GC content using R function *smooth.spline*. Next, we calculated the predicted over/under–representation of each gene from each sample based on the fitted spline, and adjusted its RPKM value to remove the effect of different GC content on mRNA level of individual genes.

### Identification of binding sites of transcription factors

TF binding data for human LCL GM12878 was from the ENCODE (Encyclopedia of DNA Elements) project;^28^ hgl9 coordinates of TF binding regions were obtained from the “Txn Factor ChIP” track (the wgEncodeRegTfbsClusteredV2.bed.gz file). We used binding regions for 50 TFs (see Table S2) that are consistently expressed in all 3 RNA-Seq datasets from Lappalainen et al.^2^ We defined a census promoter region as the 1000 bp upstream to 1000 bp downstream of gene transcription start site (TSS). Overlaps between TF binding regions and gene promoter regions were detected using GenomicRanges^23^ and gene annotations from Ensembl Genes 69.^24^

### Enrichment analysis for TF binding, TF knock down and KEGG pathways

Enrichment tests were performed by upper-tailed hypergeometric test using R function *phyper*, Bonferroni corrected for the number of tests, to determine whether binding of individual, pairs or combinations of TFs are enriched in promoter regions of CGGs, or whether KEGG pathways or genes that are differentially expressed from the knockdowns of TF genes from Cusanovich et al.,^7^ are significantly enriched in CGGs. We retrieved Ensembl gene IDs for 229 KEGG pathways^29^ using org.Hs.eg.db.^30^ To identify CGGs with mitochondrial localization, the list of 1158 human mitochondrial genes were obtained from MitoCarta2.0.^31^

### Analysis of protein-protein interactions

To test for the enrichment of protein-protein interactions (PPIs) among proteins encoded by CGGs, we used PPI data from STRING.^32^ P values for PPI enrichment were calculated based on a random background model that preserves the degree distribution of input proteins using function *get_summary* from the STRINGdb^33^ package.

### Analysis of alternative splicing

RNA-Seq reads for LCLs from 88 Yoruba and 88 Caucasian individuals^2^ were mapped to genomic regions of retained intron (RI) and skipped exon (SE) events using TopHat2.^34^ Annotations of Rl and SE events were obtained from MISO^35^ (specifically the *miso_annotations_hgl9_v2.zip* file). Only uniquely mapped reads were considered in the following analysis. To quantify the level of alternative splicing, MISO was used to detect alternative splicing events and calculate the percent spliced in (PSI) values. To test the association between expressions of spliceosome genes in the combined CGG and PSI values using YRI and CEU RNA-Seq data,^2^ we calculated the PC1-PC20 eigenvectors of expression profiles of the spliceosome genes to represent their shared expression pattern across multiple individuals using R function *svd*. Separately, we calculated the average R^2^ for the expression profile of each spliceosome gene and the splicing profile of all RI or SE events in a linear regression model. Average correlation between splicing profile of individual RI or SE events and expression profiles of spliceosome genes were calculated using Pearson’s correlation coefficient *r*.

### Data Availability

All mRNA level data used for gene expression analyses are previously published^2,18,19,25^. Other data necessary to support the conclusions of this work are represented fully within the article or in the supplemental material.

## Results

### Experimental design

Our experimental strategy is based upon using PCA to detect groups of genes whose mRNA levels are covarying. We use three published mRNA level data sets of human LCLs derived by mRNA-Seq, the first from Lappalainen et al.^2^ containing 462 individuals from 5 populations: 91 Caucasians (CEU), 89 Yoruba (YRI), 95 Finns (FIN), 94 British (GBR) and 93 Toscani (TSI); the second from Pickrell et al.^18^ containing 69 Yoruban individuals; and the third from Montgomery et al.^19^ containing 60 Caucasians. We also analyse a gene expression microarray dataset from Stranger et al.^25^ containing 109 Caucasians. Full details are in Materials and Methods.

Our approach is based upon the view that *trans* genetic influences should affect multiple genes and that we do not know the numbers of genes, nor the scale of variation, that are likely to be affected. The mRNA level of any gene is conceptually controlled by multiple *cis*-and trans-acting elements and so the mRNA level in any given individual will be set by the particular combinations of *cis* and *trans* variables that that individual contains. Each *trans* variable, such as the level of a single TF, will act in concert with many other TFs in the individual, contributing to the final level of mRNA of the cognate genes in the individual.

Based upon this view, we would expect that such expression data assayed from multiple individuals in a population should demonstrate covariation associated with these shared, *trans*, genetic regulatory influences, along with contributions from both other biological regulatory influences and related technical artefacts. Such shared variation is readily captured by PCA, and building on the findings of Goldinger et al.^16^ and related works,^36,37^ we would predict that different principal components will contain differing proportion of contribution from both biological and batch influences upon of variation. Under this assumption, we build our analytical approach as follows. We can apply PCA on the mRNA level data and the first PC will capture some shared variance, which in principle is caused by a shared influence. Similarly, the second PC will capture variance that might be due to a second influence and this process can be repeated sequentially. Given we have argued that *trans* influences are associated with correlated mRNA level variation in multiple genes, we use correlation statistics to interpret the effect of removal of variance by PCA, recognising that at this stage we cannot distinguish between correlation due to *trans* or batch influences. The process we have developed is detailed in Figure 1. We first set an absolute correlation level above which we deem the correlation to be significant using a permutation approach (see below and Figure S2 for justification of correlation thresholds). To illustrate the process, let us assume we have mRNA level data on multiple genes derived from LCLs of multiple individuals. In the data prior to PCA analysis (mRNA level matrix X_1_), we calculate all pairwise mRNA level correlations (correlation matrix *C*_1_) and identify the genes whose mRNA levels are correlated better than the threshold (Figure 1A, Figure 1B). Following the removal of PC1 (yielding matrix *X*_2_) we recalculate the correlation (matrix *C*_2_) and again find genes correlated above the threshold. In Figure 1C, the Venn diagram is composed of two sets of genes: those genes that are correlated above the threshold in the source data (*C*_1_) and genes that are correlated above the threshold in the PCI removed data (*C*_2_). For the genes displayed in the sector marked PC1* in Figure 1C, we suggest that filtering out PC1 has removed variance from these genes that corresponds to shared, and possibly *trans*, influences and so reduces their correlation below threshold. We then remove PC2 (*X*_3_) and again recalculate the correlation of all pairs of genes (*C*_3_) and identify the genes that fall into the Venn diagram sector marked PC2* in Figure 1C. Again, we suggest filtering out PC2 has removed shared, possibly *trans*, variance influencing this set of genes. Thus, in each PC there are a set of genes that are better correlated than the threshold but that in the next PC no longer meet the threshold (see Figure 2). Each set of genes we define as “correlating group of genes” (CGG) whose covariation is removed by PC1 or PC2 (Figure 1C). We use a nomenclature that is based upon the covariation detectable in one set of data (*X_n_*) and removed by the next PC (*X*_*n*+1_), and we call these genes the “CGG of PC_n_” or “PC_n_ CGG”.

**Figure 1.**
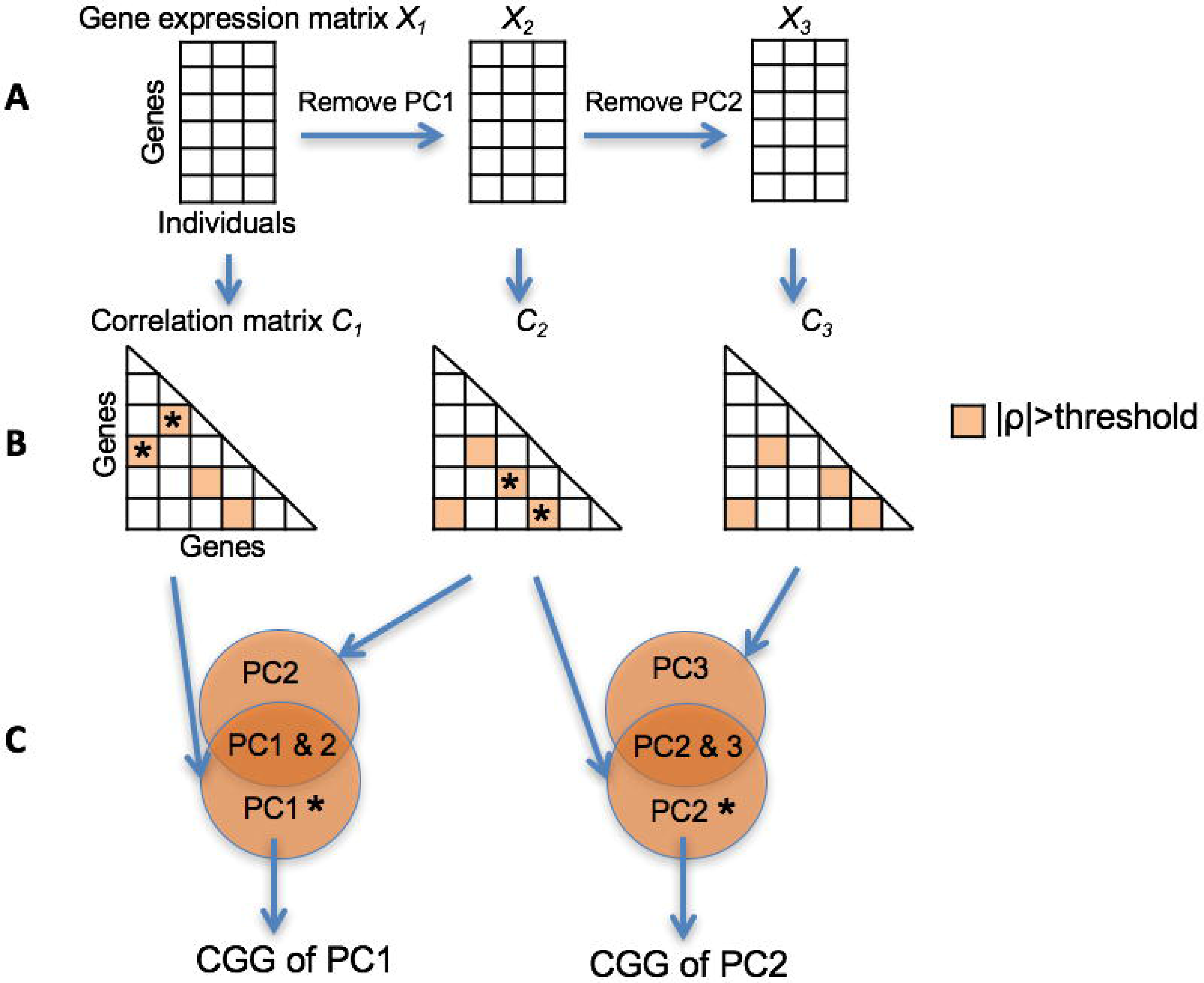
Schematic of experimental design to identify CGGs. (A) PCA is applied to a set of appropriate, pre-processed mRNA level measurements (matrix *X*_1_) derived from multiple human LCLs. PCA is used sequentially and after each principal component has been removed we generate a new mRNA level matrices *X_n_* to create 20 matrices *X*_1_–*X*_20_ (*X*_1_–*X*_3_ are shown here). (B) Pair-wise mRNA level correlations are calculated for each mRNA level matrix to yield a matrix of correlations of all possible gene pairs (matrices *C*_1_–*C*_20_, *C*_1_–*C*_3_ are shown here). Gene pairs with correlations above the threshold are highlighted in orange. (C) Correlating group of genes (CGG) of PCI are defined as genes included in gene pairs with correlations above the threshold in correlation matrix *C*_1_ but not contained in any gene pairs with correlations above the threshold in correlation matrix *C*_2_. Similarly, CGG of PC2 are defined as genes included in correlated gene pairs from *C*_2_ but not in correlated gene pairs from *C*_3_.

**Figure 2.**
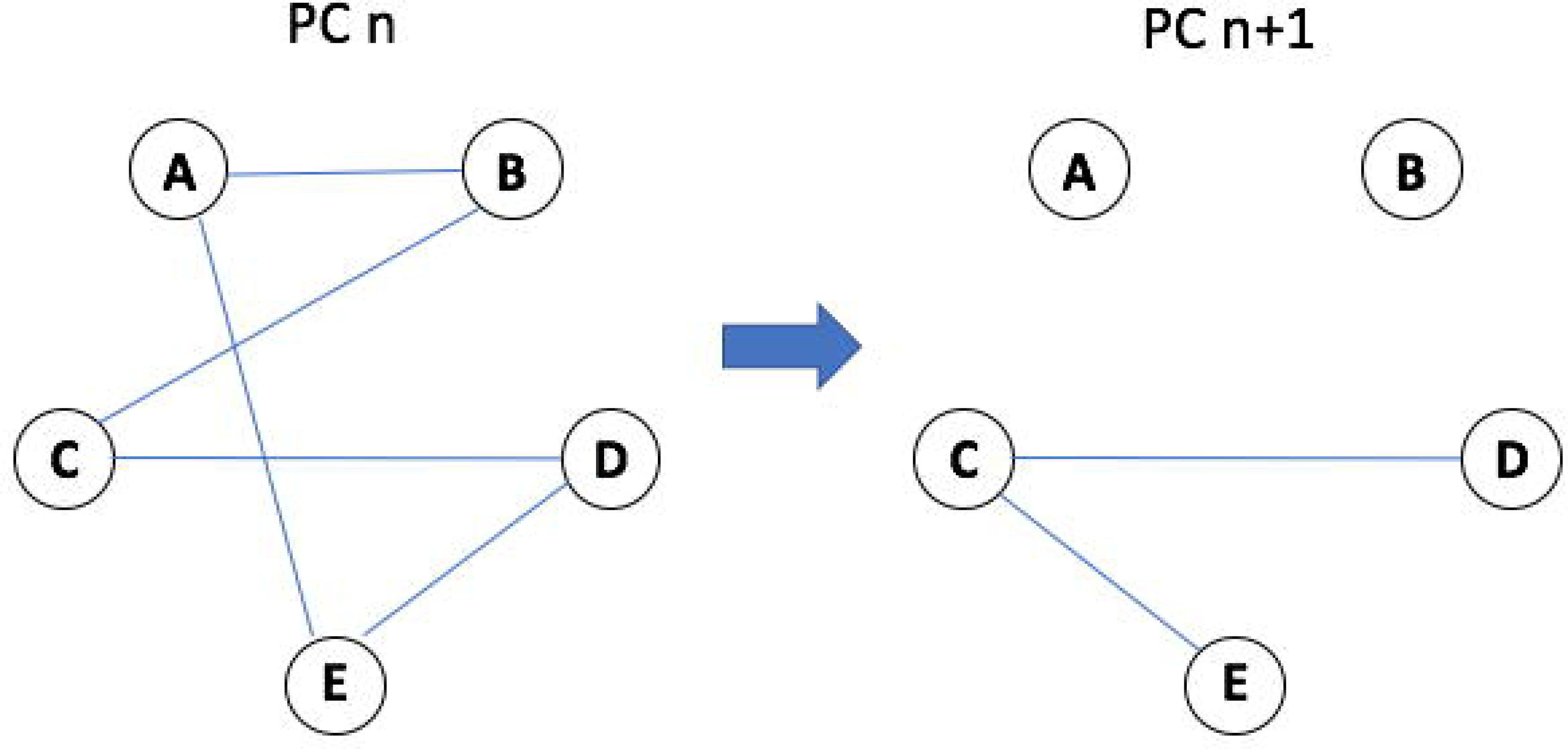
Example of correlating groups of genes. In PC_n_ gene expression data, 5 different genes A–E are connected with each other by edges representing correlation coefficient above the threshold (blue lines). Correlations for gene pairs below the threshold are not shown. In PC_n+1_ data, all correlations for A and B fall below the threshold, indicating that they are no longer correlated with any other gene. So, gene A and B are defined as CGG of PC_n_. Please note that two correlation edges for gene E are lost in PC_n+1_, but a new edge between gene C and E is formed. Therefore, gene E does not match the definition of CGG.

At this stage, we cannot determine if the covariation that defines our CGGs is either due to a batch effect, shared variation of biological origin, including *trans* influences, or some combination of the two. To determine if batch effects can, or cannot, best explain the behaviour of the CGGs, we designed several analyses to *1)* carry out multiple replication studies to show that it is implausible that the effects we observe are due to experimental artefacts, and *2)* employ multiple analyses of different biological properties of the CGGs to show that the distribution of these properties is also incompatible with experimental batch effects.

Replication analyses are carried out using the same procedure on multiple datasets and comparing the overlap of CGGs identified by each dataset. We use 3 classes of analysis (Table 1):

1. Within and between population replication.
2. Technical replication by analysis of the same mRNA samples with library generation and sequencing in 2 independent labs.
3. Cross platform replication using mRNA samples of the same individuals analysed using different analysis techniques.

**Table 1.**
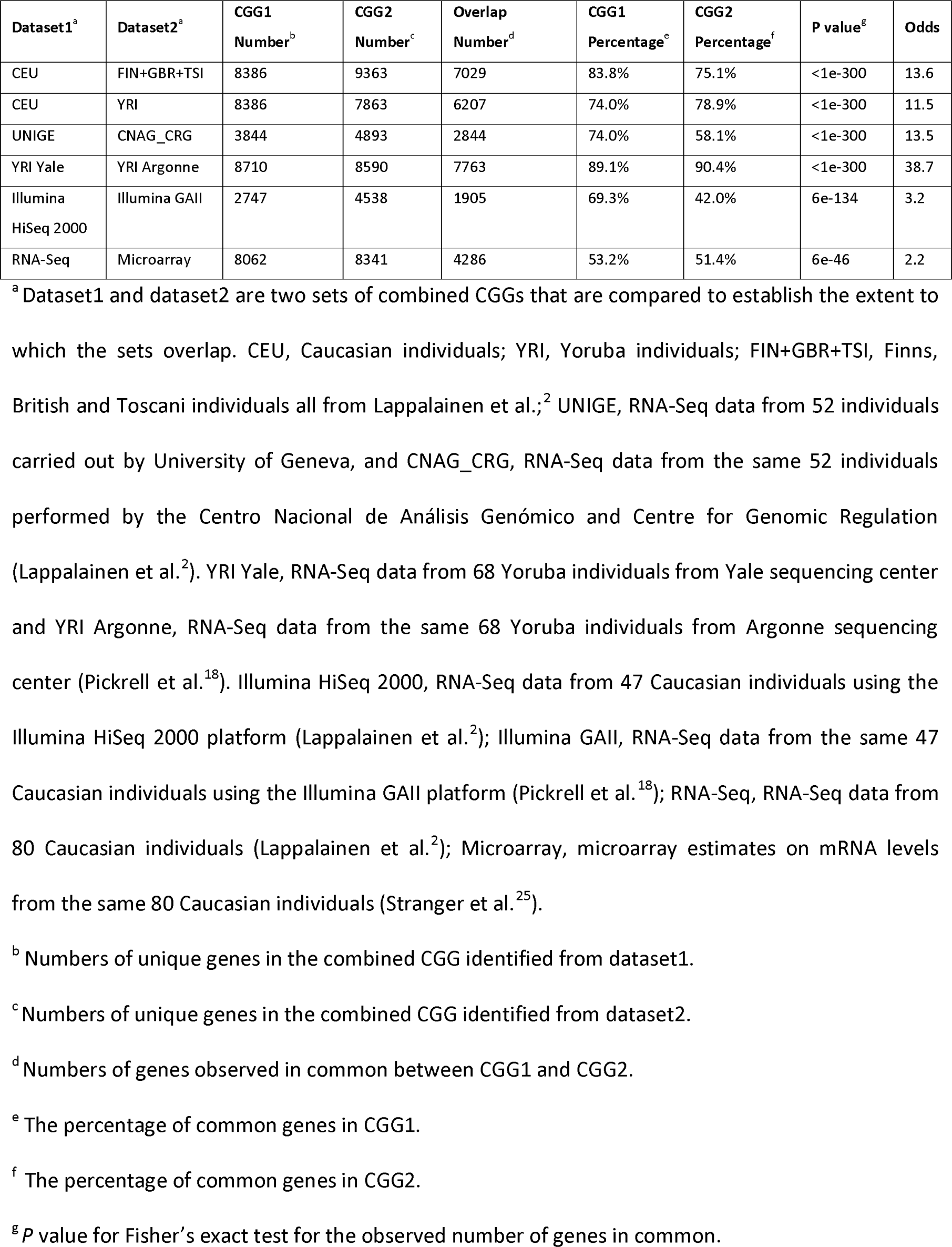
Overlap of CGGs identified from different datasets.

Our reasoning is that technical batch effects affecting CGGs are less likely to be replicable between laboratories and across different platforms, but that CGGs should replicate within and between different populations.

We then ask if the CGGs hold biological properties that would not logically be observed in sets of genes that are simply generated as a consequence of underlying batch effects. We use four approaches: *1)* we test for overabundance of TFs bound to the census promoter region of CGGs, as binding might be expected to be enriched in the case of *trans* influence rather than batch effects; *2)* we test for overabundance of genes that are sensitive to the knockdown of individual TFs, which are expected to be enriched in CGGs due to *trans* influence of TFs; *3)* we test for overabundance of functional descriptors (KEGG pathways^38^) in the CGGs: as gene groupings artefactually formed by batch effects are expected to be independent of any annotated biological function; and finally *4)* we look at the protein-protein interactions (PPIs) of the proteins encoded by CGGs and ask if they interact more than random expectation: again, batch effects are not expected similarly to influence genes that encode proteins with more interactions than random expectation. The same approaches have been used previously in related analyses^7,10,12^ to support a biological rather than batch origin of changes in gene expression.

Finally, we show that inter-individual variation in expression of a set of genes involved in mRNA splicing and also contained in CGGs is associated with variation in splicing patterns within the same individuals, suggesting that the shared variation of mRNA level in CGGs may have phenotypic consequences.

These multiple lines of evidence lead to the conclusion that mRNA levels of substantial numbers of genes are influenced collectively in *trans*, and that their covariation has phenotypic effects upon at least one property of human cells.

### Defining correlation thresholds

We set a correlation threshold for our analyses by systematically permuting the individual labels of each gene expression profile for each mRNA level matrix created by sequential removal of PCs. We reason that permutation reveals background noise, and therefore any correlations observed in unpermuted data are likely to derive from either biological or background noise. We applied 1000 permutations and recorded the maximum absolute correlation for each mRNA level matrix as the null distribution. In Figure S2 we graph for each PC removal step the correlation values corresponding to *P*=0.001, *P*=0.01 and *P*=0.05 based on the null distribution of correlation generated from 1000 permutations. We identify correlation thresholds for all sets of mRNA level matrices used in our analyses (Table S1); for the Lappalainen et al.^2^ data sets this is an absolute correlation more than 0.64. We examined the number of gene pairs with correlation above the threshold in mRNA level data with sequential removal of PCs (Figure S3A), and observe that it decreases until about PC20, plateaus until about PC40 then steadily increase. We also observe (Figure S3B) that in randomly permuted mRNA level data there are only few gene pairs correlating above the threshold in PC1–20, followed by a steady increase of numbers of correlating gene pairs with subsequent removal of PCs. This observation suggests that most biological effects influencing gene pair correlations should be within the first 20 PCs. We therefore limit our analyses to the first 20 PCs.

### Defining correlating groups of genes

To identify potential genes under *trans* regulation we used the RNA-Seq data for 88 individuals each drawn from the Caucasian, Yoruban and combined Finns, British and Toscani population from Lappalainen et al.,^2^ and carried out the process shown in Figure 1 to identify CGGs. In Figure 3 we show the number of genes within 20 sets of CGGs as consecutive PCs are removed (54–2137, 75–2168 and 75–2155 genes for Caucasian, Yoruban and combined Finns, British and Toscani population, respectively). The pattern of above threshold correlations of genes can be very complex, and so the gene-pair correlations are conveniently treated as a network with each gene as a node connected by edges representing correlations above the threshold. Summary statistics of these networks from each population are in Table S3, and as an example of the complexity of the correlations, the gene *PPIAP29* identified in PC1 CGG of combined Finns, British and Toscani population from the Lappalainen et al.^2^ has 997 pairwise absolute correlations better than 0.64 — the highest number of connections in our data. An example correlation network (PC7 CGG from RNA-Seq of Caucasian individuals from Lappalainen et al.^2^) is shown in Figure S4 which is a network defined by 1578 genes and 6268 gene pairs with |ρ| >0.64. This network contains around 0.5% of the 1,244,253 total possible gene pairs definable between member genes and 56% of all correlations are poor with absolute correlation level between −0.2 and 0.2 (Figure S4A). The network is non-homogeneous and has obvious multiple clusters (Figure S4B). Generally, the correlations of gene pairs in CGGs correspond to 0.3–1.4% of all gene pairs in the mRNA level data with removed PCs (Table S3).

**Figure 3.**
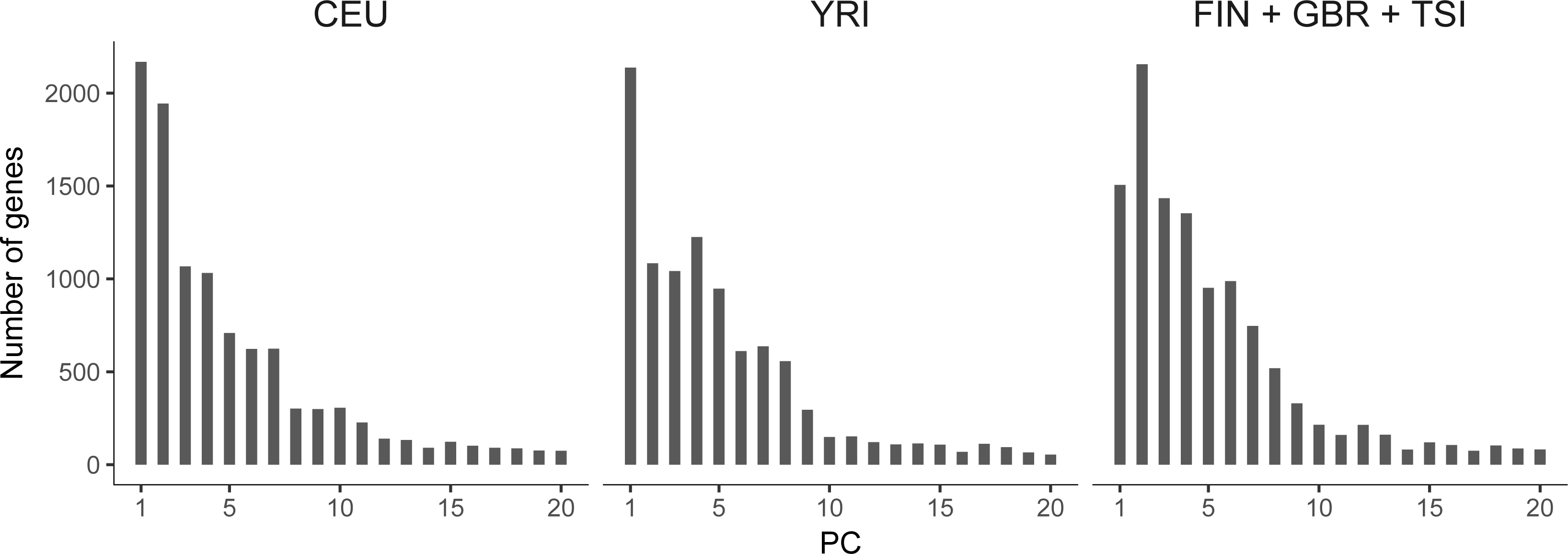
Numbers of genes in CGGs defined by principal components. The x-axis is sequential PCs that are removed from mRNA level data to identify CGGs, and the y-axis is numbers of genes identified as CGGs defined by the relevant PC. Left panel, CEU: Caucasian individuals; center panel, YRI: Yoruba individuals; right panel, FIN+GBR+TSI: Finns, British and Toscani mixed individuals. mRNA level data is from Lappalainen et al.^2^

We tested the behaviour of our approach by conducting simulations that are detailed in Materials and Methods and designed to study the behaviour of sets of genes that have artificially offset expression variation that will induce correlations and that may therefore be expected to define CGGs in our analyses. In brief, we used the RNA-Seq data of 88 Yoruban individuals taken from Lappalainen et al.^2^ with individual labels randomly permuted on a per gene basis. We hypothesized that the number of covarying genes and the scale of offset would both influence the detection of CGGs, so we randomly selected 4 groups of genes containing 200, 500, 1000 and 2000 genes, and adjusted the normalised expression levels of genes in each group with an added offset of 0.5, 1.0, 1.5 or 2.0 standard deviations, corresponding respectively to 1.15, 1.33, 1.54 and 1.78 fold mRNA level increase in random 44 of the 88 individuals. The 1.78 fold adjustment is not unrealistically large as in the 88 individuals there is an average of 2.4 fold change between individuals, and following the 1.78 fold adjustment, 80% of the adjusted gene’s mRNA level still fall within the extremes of the unadjusted expression values. We then attempted to identify which genes were classified as being in CGGs. We repeated this analysis 100 times using different random selections of genes and below we use the relevant median values of the 100 analyses. The 1.15 and 1.33 fold adjustment for all 4 groups of genes results in not more than 1% of the genes being defined as CGGs. In contrast, the 1.54 fold adjustment results in 10%, 19%, 29.2% and 40.6% of, respectively, the 200, 500, 1000 and 2000 genes being identified in the first PC as CGGs. For the 1.78 fold adjustment, the equivalent figures are 93.2%, 97.6%, 99.1% and 99.7%. False positives were not detected in any set of genes, and false negatives (failure to identify a gene with an offset as being a member of a CGG) are decreasing with the increase of gene numbers and offset size (Table S4A). These data suggest that there are two determinants of the efficiency of detection of CGG: the number of genes and the extent of variation driving the correlation.

Covariation with a biological origin should be replicable but these analyses will themselves by confounded by batch effects. We would expect batch effects could have two extreme influences upon the replication of CGGs: they could either disrupt covariation such that individual genes were no longer detectable as CGGs in two or more populations, or they could alter patterns of covariation such that the PCs that detected the same CGGs were different in the two samples. We therefore carried out a second simulation to address how batch effects might interfere with detection of covariation in our analyses. We used the unpermuted Yoruban data (Figure 3) where we had previously detected 75–2168 CGGs but we randomly selected 4 groups of genes containing 200, 500, 1000 and 2000 non-CGG genes (defined as all expressed genes other than genes in any CGG), and adjusted the expression of genes in each group with on average 1.15, 1.33, 1.54 and 1.78 fold increase in random 44 of the 88 individuals. This adjustment can be treated as a batch effect by comparing the recovery on a PC by PC basis of the CGGs in the presence or absence of the gene expression adjustment. We identified the union of genes that are identified as being a member of CGG defined by any PC up to 20 PCs; this list is the "combined CGG". We repeated this analysis 100 times and calculated the relevant median values. Recovery of CGG in any PC in the presence or absence of the batch effects ranges from 97.7% to 99.8%, but importantly as both the number of genes and the offset becomes larger, the PC that defines the CGG in the original data compared to the batch effects containing data changes. For the 2000 gene set, 7.72%, 21.8%, 42.9% and 57.4% of combined CGG respectively for the 1.15, 1.33, 1.54 and 1.78 fold offset of gene expressions, are identified as CGG members in a different PC (Table S4B). This analysis tells us that replication of CGGs with artificial batch effects will often not be based upon the same genes being defined by the same PC; instead, we define replication as the same genes being defined in the combined CGG without considering from which PC they were detected.

Based upon the simulations, we use the overlap of combined CGG as a criterion of replication. We identified the combined CGG for RNA-Seq data of 88 individuals each drawn from the Caucasian, Yoruban and combined Finns, British and Toscani population from Lappalainen et al.,^2^ and in Table 1 we show the overlap of genes within the combined CGG detected within a population: the combined CGG from 88 Caucasian compared to the combined CGG from 88 Finns, British and Toscani (Lappalainen et al.^2^) has 84% and 75% overlap with an odds ratio of 13.6 (Table 1, row 1); the CGGs from Caucasian and Yoruban has 74% and 79% overlap with an odds ratio of 11.5 (Table 1, row 2). In Figure S5, we show replication analyses for the CGGs defined by PC1-10, and note that there is significant replication of CGGs but the best overlaps are not always of CGGs defined by the same PC in the two analyses.

### Technical replication of CGGs

To test for technical replication of combined CGG between experiments, we used samples from Lappalainen et al.^2^ who report the mRNA-Seq analysis of the same 52 mRNA samples with library generation and sequencing in 2 independent laboratories, and we observe replication with 74% and 58% overlap and an odds ratio of 13.5 (Table 1, row 3). Similarly, 68 Yoruban mRNA samples were sequenced by 2 independent laboratories reported by Pickrell et al.,^18^ and we observe replication with 89% and 90% overlap and an odds ratio of 38.7 (Table 1, row 4).

To test for replication of combined CGG across machine types, we compared combined CGG of the same individuals analysed in different labs using different analysis machines or techniques. We compared combined CGG from Illumina GAII and HiSeq2000 analysis of the same 47 individuals analysed by Pickrell et al.^18^ and Lappalainen et al.,^2^ respectively (Table 1, row 5). Based upon Fisher’s exact test the genes in combined CGG replicate across the analyses with 69% and 42% overlap and an odds ratio of 3.2. We also compared combined CGG from the same 80 individual’s mRNA analysed on microarrays and Illumina HiSeq2000 by Stranger et al.^25^ and Lappalainen et al.,^2^ respectively (Table 1, row 6), and observed 51% and 53% overlap with an odds ratio of 2.2, which is for data derived from two very different methodologies.

In Figure S6, we show the replication of CGGs defined by PC1-10 from RNA-Seq data of different labs, and from gene expression data of different machine types in Figure S7. The best overlaps of CGGs are not from the same PC suggests that different pattern of technical batch effects between experiments may influence the distribution of CGGs across multiple PCs. The list of CGGs for each dataset mentioned above are provided in Table S5, and the IDs of individuals for each dataset analysed are in Table S6.

### Batch effects

The fact that we see replication of CGGs across multiple analyses, populations and platforms makes it difficult to suggest that experimental batch effects alone are the source of the shared behaviour of gene expression. Later we will show that the genes contained within the CGGs indeed have a number of biological properties that cannot be accounted for by experimental variations but we accept that batch effects must have an influence upon overall correlation of mRNA levels. We therefore repeated our analyses but first used two methods commonly used to remove technical artefacts. PEER (probabilistic estimation of expression residuals) from Stegle et al.^27^ is a software that is widely used for removing batch effects from mRNA-Seq data. PEER removes 61.7%, 63.0% and 64% of total variance of the mRNA levels of LCLs from Yoruban, Caucasian, and the Finns, British and Toscani from Lappalainen et al.,^2^ and we identify 651, 693 and 644 combined CGG, compared with 7863, 8386 and 9363 combined CGG identified without PEER, with an overlap of 6.1%, 6.2% and 5.4% of genes, respectively. PEER is very effective at reducing variance of gene expression, and so it is unsurprising that this leads to a very substantial drop in CGG numbers, but the analysis is likely to be removing both technical and biological sources of variation.

In the second case we used an approach that is based upon mRNA level variation that was due to GC base composition biases in mRNA-Seq (Materials and methods). This method was developed by Pickrell et al.^18^ to identify sample to sample deviation from expectation of mRNA-Seq read counts mapped to exons separated by GC base composition and to correct for such deviation. For the Yoruban mRNA level data^2^ with GC bias correction we obtained 6762 genes in the combined CGG following the same data analysis procedure. Comparing with the combined CGG without GC bias correction, the recovery rate of CGG is 93% with 61% genes shifted among their original defining PCs, which suggests that GC bias is not a major explanation for correlated behaviors of CGG.

In both cases these methods are designed to leave an expression level of an individual gene as unaffected as possible by shared influences; in reality we believe it is most probable that removal of shared variations or correction of deviation from random or other expectation is likely to be removing both technical artefacts and biological signal from some genes and, inevitably, as greater variance is removed by filtering out initial PCs, correlation in mRNA levels will necessarily decrease to a minimum (Figure S3A). As a consequence, we rely on our PCA approach to remove variance systematically, recognizing that the sources of this variance cannot be established from identification of correlations alone.

Collectively our analyses support the view that the behaviour of CGGs is unlikely to be best explained by batch effects as we see replication across both platforms and laboratories, individuals and populations. We therefore set out to examine if CGGs have biological properties that are not associated with the technology of mRNA-Seq at a frequency greater than random expectation: we argue that association of such biological properties with CGGs is supportive evidence for a biological explanation for correlations in mRNA levels.

### Some TFs show over abundant binding to census promoter regions of CGGs

Allocco et al.^39^ have shown that the expression level of genes that share TFs bound to their cognate promoter are better correlated than those that share fewer. We therefore analysed the TF binding in the 1000 bp up-and down-stream of the TSS of sets of the CGGs reasoning that if the behaviour of CGGs is indeed of biological origin we might expect particular TFs to be more associated with CGGs than random expectation. We use the binding site data from 50 TFs (listed in Table S2) whose binding has been established in a lymphoblastoid cell line by the ENCODE project^28^ and use hypergeometric test, with appropriate correction for multiple tests, to identify enrichment for binding of any TF to the cognate promoter regions of the combined CGG using RNA-Seq data of 3 populations containing 88 Yoruban, 88 Caucasian and 88 individuals from the British, Finns, Toscani populations (data of Lappalainen et al.).^2^ These results are displayed in Table 2. 38 of the 50 TFs have significantly enriched binding to the promoter regions of genes in the combined CGG defined from one or more populations, and in Table S7 we show the enrichment within the CGGs identified from individual PCs. In Figure S8 we show that there is substantial similarity in overabundance of TF binding to the combined CGG of 3 populations.

**Table 2.** Enrichment of individual transcription factors bound to promoter regions of CGGs.

| TF symbol | CEU |  | YRI |  | FIN+GBR+TSI |  |
| --- | --- | --- | --- | --- | --- | --- |
|  | N | P<sup>a</sup> | N | P<sup>a</sup> | N | P<sup>a</sup> |
| ATF3 | 826 | 1.8e-05* | 782 | 1.0e-07* | 893 | 2.0e-05* |
| BCLAF1 | 1044 | 3.7e-05* | 974 | 1.7e-05* | 1131 | 2.0e-05* |
| BRCA1 | 222 | 1.7e-03 | 216 | 1.3e-05* | 236 | 1.4e-02 |
| CHD2 | 1691 | 1.5e-10* | 1615 | 1.0e-17* | 1827 | 3.6e-12* |
| EBF1 | 2164 | 3.3e-04* | 2067 | 5.5e-07* | 2428 | 2.5e-09* |
| EGR1 | 3425 | 1.4e-08* | 3187 | 1.7e-10* | 3776 | 9.2e-16* |
| ELF1 | 5114 | 5.3e-16* | 4833 | 5.0e-29* | 5595 | 8.9e-23* |
| ETS1 | 1344 | 1.4e-08* | 1271 | 9.3e-12* | 1472 | 1.5e-11* |
| FOS | 866 | 4.3e-03 | 833 | 3.2e-05* | 916 | 1.5e-02 |
| GABPA | 2468 | 2.7e-09* | 2343 | 8.7e-16* | 2683 | 3.7e-12* |
| IRF4 | 938 | 3.3e-05* | 888 | 4.0e-07* | 1025 | 6.3e-06* |
| MAX | 501 | 1.3e-01 | 493 | 8.0e-04* | 548 | 1.5e-01 |
| MEF2A | 1060 | 1.0e-04* | 1017 | 5.2e-08* | 1163 | 2.9e-05* |
| MEF2C | 456 | 2.1e-03 | 432 | 4.7e-04* | 497 | 3.5e-03 |
| MYC | 1176 | 4.8e-13* | 1111 | 1.8e-15* | 1260 | 6.5e-12* |
| NR2C2 | 345 | 4.2e-04* | 328 | 7.8e-05* | 355 | 2.1e-02 |
| NRF1 | 2394 | 3.0e-07* | 2233 | 1.2e-09* | 2604 | 1.4e-09* |
| PAX5 | 2680 | 8.8e-08* | 2508 | 2.2e-09* | 2923 | 2.0e-09* |
| PBX3 | 1324 | 1.2e-06* | 1249 | 8.0e-08* | 1401 | 5.9e-04* |
| POU2F2 | 2684 | 7.0e-20* | 2539 | 3.6e-27* | 2884 | 1.3e-17* |
| RELA | 750 | 1.9e-06* | 702 | 1.2e-07* | 799 | 8.9e-05* |
| RFX5 | 1038 | 3.7e-03 | 997 | 3.3e-07* | 1146 | 5.4e-06* |
| RXRA | 632 | 2.0e-06* | 599 | 1.9e-07* | 677 | 1.4e-05* |
| SIN3A | 4057 | 1.4e-18* | 3811 | 7.0e-26* | 4370 | 4.6e-19* |
| SIX5 | 2137 | 5.3e-14* | 2056 | 6.2e-23* | 2301 | 2.1e-13* |
| SP1 | 3426 | 7.4e-14* | 3277 | 1.9e-25* | 3716 | 6.8e-16* |
| SPI1 | 1976 | 1.8e-05* | 1873 | 2.8e-09* | 2201 | 5.9e-09* |
| SRF | 1775 | 7.7e-10* | 1658 | 6.4e-12* | 1896 | 6.0e-08* |
| TAF1 | 2635 | 7.5e-31* | 2446 | 8.5e-31* | 2797 | 2.0e-25* |
| TBP | 2886 | 2.7e-26* | 2692 | 1.2e-31* | 3101 | 7.8e-26* |
| TCF12 | 2083 | 1.2e-10* | 1946 | 2.5e-12* | 2261 | 1.7e-11* |
| USF1 | 1604 | 4.9e-04* | 1533 | 2.3e-07* | 1765 | 1.9e-06* |
| USF2 | 1454 | 1.2e-02 | 1393 | 9.9e-06* | 1586 | 1.6e-03 |
| WRNIP1 | 344 | 3.0e-07* | 327 | 1.7e-08* | 351 | 2.1e-04* |
| YY1 | 4273 | 1.9e-19* | 4008 | 8.8e-25* | 4617 | 6.6e-20* |
| ZBTB33 | 795 | 4.2e-04* | 756 | 1.6e-05* | 852 | 1.6e-03 |
| ZEB1 | 1922 | 7.7e-03 | 1821 | 2.9e-05* | 2136 | 2.5e-05* |
| ZNF143 | 2506 | 2.4e-06* | 2421 | 3.4e-14* | 2768 | 7.4e-11* |

The table lists the TFs that are found to be significantly associated to the combined CGG in at least one population. CEU, Caucasian individuals YRI, Yoruba individuals; FIN+GBR+TSI, Finns, British and Toscani individuals all from Lappalainen et al..<sup>2</sup> N: The number of CGGs with individual transcription factors bound to promoter region defined as $\pm 1000$ base pairs of TSS; P: *P* value for hypergeometric test for enrichment of binding.
<sup>a</sup> $P < 1.0 \times 10^{-3}$ (Bonferroni corrected significance threshold) are labeled with \*.

In summary, multiple TFs compared to random expectation appear to be over abundantly associated with census promoter regions of CGGs, which have been identified from multiple populations. We recognise of course that classification of genes as having a bound TF is necessarily an inference based upon the ENCODE^28^ analysis of a single LCL; Although there is inter-individual variation in TF binding (*e.g*. 7.5% of binding regions of NFKB were found different among individuals by Kasowski et al.),^40^ the scale of the overlaps suggests that such effects would have rather limited influence upon our analysis. At the very least, we would not expect enrichment of any TF binding to cognate promoters to be associated with batch effects, adding further support to the view that there is a biological basis for CGGs’ behaviours.

### Genes known to be sensitive to single TF knockdown are enriched in CGGs

Our results suggest the possibility that the variation in the amount or activity of the appropriate TFs may be involved in generating the correlated behaviour of CGGs, but numerous lines of experimental evidence show that for most genes binding of a single TF to a cognate promoter region is neither necessary nor sufficient for controlling mRNA level.^41–43^ Consequently, our detection of overabundance of TF binding to some CGGs does not necessarily imply a causal relationship between variation in those overabundant TF bindings and variation in their target genes’ mRNA levels. Cusanovich et al.^7^ tested the relationship directly by studying the effects of knockdown of expression of 59 TFs and chromatin modifiers (TFs used in this study are listed in Table S8). They were able to identify sets of genes whose expression levels were directly influenced by knockdown of single TFs. If the relationship of CGGs with TF is causal of shared behaviour, we might expect the genes identified by Cusanovich et al.^7^ are enriched in relevant CGGs. In Table 3 we show that there is a statistically significant overlap of genes identified as combined CGG with genes defined as being sensitive to knockdown of 29 different TFs (see Table S9 for enrichment in CGGs from individual PCs). Of the 29 TFs, 9 are also analysed for binding in the ENCODE data: of these 9 TFs, 8 (IRF4, RELA, POU2F2, PAX5, SP1, TCF12, USF1, YY1) demonstrate both overabundance of binding sites, and enrichment for targets of TF knockdowns.

**Table 3.** Enrichment of differentially expressed genes from transcription factor knockdown in CGGs.

| TF Knockdown | CEU |  | YRI |  | FIN+GBR+TSI |  |
| --- | --- | --- | --- | --- | --- | --- |
|  | N | P<sup>a</sup> | N | P<sup>a</sup> | N | P<sup>a</sup> |
| *ARNTL2* | 443 | 2.0e-04* | 447 | 2.1e-09* | 501 | 3.1e-08* |
| *CEBPG* | 269 | 2.0e-02 | 275 | 2.0e-06* | 286 | 2.2e-02 |
| *CEBPZ* | 263 | 1.4e-03 | 238 | 6.4e-03 | 287 | 7.3e-04* |
| *CREBBP* | 1516 | 2.7e-08* | 1416 | 1.6e-09* | 1624 | 6.7e-07* |
| *DIP2B* | 508 | 7.5e-04* | 470 | 3.8e-04* | 548 | 3.7e-04* |
| *E2F1* | 270 | 8.2e-04* | 244 | 4.1e-03 | 285 | 3.4e-03 |
| *ESRRA* | 375 | 1.5e-06* | 333 | 2.5e-04* | 373 | 3.8e-02 |
| *HOXB7* | 506 | 6.9e-04* | 466 | 1.2e-03 | 531 | 8.6e-03 |
| *IRF4* | 2203 | 4.7e-12* | 2062 | 1.3e-14* | 2371 | 1.4e-10* |
| *IRF5* | 546 | 7.2e-05* | 517 | 6.9e-06* | 566 | 1.3e-02 |
| *NFE2L1* | 525 | 2.5e-07* | 473 | 1.9e-05* | 528 | 1.1e-02 |
| *NFKB2* | 663 | 2.1e-04* | 606 | 2.4e-03 | 703 | 4.7e-03 |
| *NR1D2* | 362 | 7.5e-04* | 328 | 6.0e-03 | 376 | 9.9e-03 |
| *NR2F6* | 694 | 4.8e-06* | 624 | 1.8e-03 | 762 | 4.5e-08* |
| *NR3C1* | 281 | 9.1e-04* | 272 | 3.6e-06* | 303 | 8.0e-04* |
| *PAX5* | 1380 | 2.2e-06* | 1292 | 1.1e-07* | 1469 | 1.2e-04* |
| *POU2F1* | 411 | 8.4e-06* | 385 | 9.2e-07* | 443 | 2.4e-06* |
| *POU2F2* | 478 | 1.1e-02 | 453 | 3.6e-04* | 536 | 6.3e-05* |
| *RAD21* | 1539 | 1.4e-05* | 1430 | 1.8e-05* | 1667 | 7.7e-05* |
| *RELA* | 240 | 5.0e-02 | 236 | 5.4e-04* | 250 | 1.5e-01 |
| *SP1* | 2023 | 2.0e-05* | 1952 | 1.7e-13* | 2212 | 4.2e-07* |
| *STAT2* | 52 | 3.3e-04* | 40 | 1.2e-01 | 50 | 6.1e-03 |
| *TCF12* | 537 | 5.0e-04* | 494 | 2.3e-03 | 577 | 1.3e-03 |
| *TFDP1* | 694 | 5.4e-07* | 647 | 1.1e-06* | 729 | 8.3e-05* |
| *TFDP2* | 468 | 6.0e-06* | 442 | 7.3e-08* | 488 | 5.8e-04* |
| *TFE3* | 427 | 1.7e-03 | 401 | 2.4e-04* | 462 | 9.1e-04* |
| *USF1* | 310 | 5.1e-03 | 297 | 9.2e-04* | 327 | 2.8e-02 |
| *YY1* | 1025 | 2.2e-02 | 957 | 6.0e-03 | 1135 | 8.9e-05* |
| *ZHX2* | 298 | 4.4e-04* | 269 | 5.7e-03 | 311 | 1.8e-03 |

The table presents genes sensitive to knockdown of individual TFs from Cusanovich et al.<sup>7</sup> that are significantly enriched in the combined CGG from at least one population. CEU, Caucasian individuals; YRI, Yoruba individuals; FIN+GBR+TSI, Finns, British and Toscani individuals all from Lappalainen et al.<sup>2</sup> N: The number of CGGs that overlap with genes exhibiting differential expression following TF knockdown by Cusanovich et al.<sup>7</sup> P: P value of hypergeometric test.
<sup>a</sup> $P < 9.6 \times 10^{-4}$ (Bonferroni corrected significance threshold) are labeled with \*.

### Combinations of TFs are overabundant in promoter regions of CGGs

Despite these analyses, it is unlikely that single TF binding is the only likely biological contribution to the variable expression of the CGGs. As discussed earlier, multiple TFs generally bind to the census promoter region of genes^28,44^ and it would therefore be useful to test for combinations of TFs binding to the census promoter. A systematic analysis of all possible combinations of the 50 TFs in the ENCODE data set is impossible due to the huge numbers of combinatorial possibilities. To overcome this, we used two approaches. Wang et al.^44^ show that 20 pairs of TFs co-bind in the genome of a human LCL: we can ask if these pairs are more commonly found in promoter regions of CGGs than would be expected at random. In the second approach, we remove PCs sequentially until we have reached a minimum number of correlated gene pairs and then search for shared binding of TFs in promoter regions of this minimal gene set; we reason that a correlated expression in just a pair of genes is the most minimal evidence of *trans* activity and therefore the shared TFs might be important in mediating *trans* influences. In both cases we use the hypergeometric test to detect overabundance of binding of combinations of TFs, ignoring their relative location, to the cognate promoter region of the combined CGG of the Yoruban, the Caucasian and the British, Finns, Toscani populations from Lappalainen et al.^2^

In the first approach, of the 20 TF pairs identified in Wang et al.,^44^ 14 are enriched in the combined CGG in one or more populations of which 8 pairs are enriched in CGGs from all three populations (Table 4). Table S10 shows the enrichment for CGGs from individual PCs. In the second approach, we sequentially removed PCs from the Lappalainen et al.^2^ Yoruban, Caucasian and British, Finns, Toscani data sets. Respectively removal of 30, 31 and 31 PCs yielded a minimum of 2142,1892 and 1719 gene pairs with absolute correlation better than 0.64 (Figure S3) of which 303, 178 and 258 had between 2 to 25 TF binding sites in common. 51 combinations of TFs were common to all 3 populations, and so we then tested for significance of overabundance of these combinations of bound TFs in the promoter regions of combined CGG and CGGs from individual PCs of the 3 populations. In Table S11 we identify significant enrichments of combinations of TFs within both the combined CGG and CGGs from individual PCs. In the combined CGG, 23 of the 51 combinations of TFs are enriched, 17 in all 3 populations, 1 in both Yoruban and British, Finns and Toscani and 5 in just the Yoruban. These data suggest that combinations of TFs might be contributing to the correlated variation in expression of CGGs and also again reinforce the view that these biological properties are not those expected of technical batch effects.

**Table 4.** Enrichment of pairs of transcription factors bound to promoter regions of CGGs.

| TF Pair | CEU |  | YRI |  | FIN+GBR+TSI |  |
| --- | --- | --- | --- | --- | --- | --- |
|  | N | P<sup>a</sup> | N | P<sup>a</sup> | N | P<sup>a</sup> |
| ATF3, NRF1 | 402 | 2.0e-03* | 383 | 4.0e-05* | 454 | 5.5e-06* |
| ELF1, YY1 | 3860 | 1.1e-19* | 3636 | 3.5e-27* | 4176 | 3.4e-21* |
| GABPA, YY1 | 2010 | 2.2e-10* | 1903 | 9.6e-15* | 2178 | 7.3e-12* |
| IRF4, JUND | 20 | 2.8e-01 | 16 | 4.7e-01 | 22 | 4.2e-01 |
| IRF4, MAX | 130 | 1.1e-01 | 118 | 2.1e-01 | 136 | 4.5e-01 |
| IRF4, MEF2A | 441 | 3.5e-03 | 421 | 1.7e-04* | 478 | 1.6e-02 |
| IRF4, MEF2C | 231 | 5.3e-02 | 215 | 3.4e-02 | 246 | 1.7e-01 |
| IRF4, RELA | 271 | 3.9e-03 | 254 | 4.6e-04* | 293 | 6.9e-03 |
| IRF4, TCF12 | 510 | 6.0e-04* | 463 | 3.1e-03 | 546 | 3.5e-03 |
| JUND, MEF2C | 9 | 5.9e-01 | 10 | 2.1e-01 | 9 | 8.9e-01 |
| JUND, SPI1 | 11 | 4.2e-01 | 11 | 8.8e-02 | 13 | 3.6e-01 |
| MAX, SPI1 | 193 | 2.1e-01 | 187 | 5.6e-02 | 211 | 2.1e-01 |
| MAX, USF1 | 354 | 1.3e-01 | 352 | 7.2e-04* | 387 | 9.7e-02 |
| MAX, USF2 | 361 | 2.0e-01 | 358 | 1.6e-03* | 396 | 1.3e-01 |
| MEF2A, SPI1 | 466 | 2.8e-04* | 439 | 2.1e-05* | 513 | 2.5e-05* |
| MEF2C, SPI1 | 186 | 9.9e-03 | 181 | 2.9e-04* | 204 | 5.4e-03 |
| PBX3, SP1 | 1072 | 1.4e-05* | 1018 | 1.2e-07* | 1129 | 1.5e-03* |
| RELA, SPI1 | 394 | 2.8e-05* | 357 | 1.1e-04* | 416 | 2.5e-03* |
| SPI1, TCF12 | 793 | 1.5e-05* | 730 | 2.8e-05* | 885 | 7.7e-09* |
| USF1, YY1 | 1186 | 1.1e-04* | 1140 | 3.9e-09* | 1301 | 6.1e-07* |

The table lists the pairs of TFs taken from Wang et al.<sup>44</sup> that are found to be significantly associated to the combined CGG in at least one population. CEU, Caucasian individuals YRI, Yoruba individuals; FIN+GBR+TSI: Finns, British and Toscani individuals all from Lappalainen et al..<sup>2</sup> N: The number of CGGs with individual pairs of transcription factors bound to promoter region defined as $\pm 1000$ base pairs of transcription start site; P: P value for hypergeometric test for enrichment of bindings.
<sup>a</sup> $P < 2.5 \times 10^{-3}$ (Bonferroni corrected significance threshold) are labeled with \*.

### CGGs encode proteins with related functions

Our data collectively suggest groups of genes are covarying in humans under complex *trans* control. We wish to understand if these genes might be associated with biological functions firstly to further test the view that the observed covariation has a biological origin and secondly to start to understand what might be, if any, the phenotypic consequences of *trans* variation. We wish to ask two questions as to the biological properties of proteins encoded in both combined and individual groups of CGGs. Firstly, do CGGs encode proteins that physically interact more commonly than random expectation, and secondly does this indicate a possible functional interaction that can be detected by enrichment of functional annotation within the KEGG pathway database?

We tested for enrichment of physical interactions of encoded proteins within the combined CGG identified from Yoruban, the Caucasian and the British, Finns and Toscani populations from Lappalainen et al.^2^ using protein-protein interactions from the STRING database.^32^ We compared the number of PPIs seen in the various CGGs with a random background model (see Materials and Methods). These analyses show that the combined CGG from any one of the 3 populations encode proteins that interact more commonly than expected by chance; the Yoruban set contains 6,008 proteins with 624,213 PPIs (*P* < 1×10^−16^); the Caucasian set of 6,389 proteins with 711,122 PPIs (*P* < 1×10^−16^) and the Finns, British, Toscani with 7,066 proteins with 831,191 PPIs (*P* < 1×10^−16^).

In Table S12 we show similar analysis for CGGs from individual PCs indicating that for all three populations there are multiple significant enrichments for PPIs within CGGs identified from individual PCs. The finding that the proteins encoded by CGGs are more likely to interact with each other than expectation again suggests that the variation in mRNA levels is unlikely to be due to batch effects within sample analyses. More interestingly in many cases proteins that physically interact have been shown to be contained within pathways of biological activity^45^ and suggests therefore that CGGs might be associated with specific pathways of biological activity.

We therefore tested for this by asking if pathways defined within the KEGG database are enriched in CGGs. In Table 5 we show the results of enrichment analysis of KEGG function terms in the combined CGG using hypergeometric test (see Materials and Methods), and in Table S13 we present the results of a similar analysis for CGGs from individual PCs. We see enrichment for 10 functions in the combined CGG: oxidative phosphorylation, pyrimidine metabolism, ribosome, spliceosome, proteasome, cell cycle, protein processing in endoplasmic reticulum, Alzheimer’s disease (AD), Parkinson’s disease (PD) and Huntington’s disease (HD). In these pathways 31–109 genes are shared in one population (Table 5) and 30–86 genes are shared within all 3 populations (Figure S9). These results suggest that, in humans there are significant numbers of genes that are functionally related and covarying, which opens the possibility that *trans* variation in genes involved in the same pathway might have phenotypic outcomes.

**Table 5.**
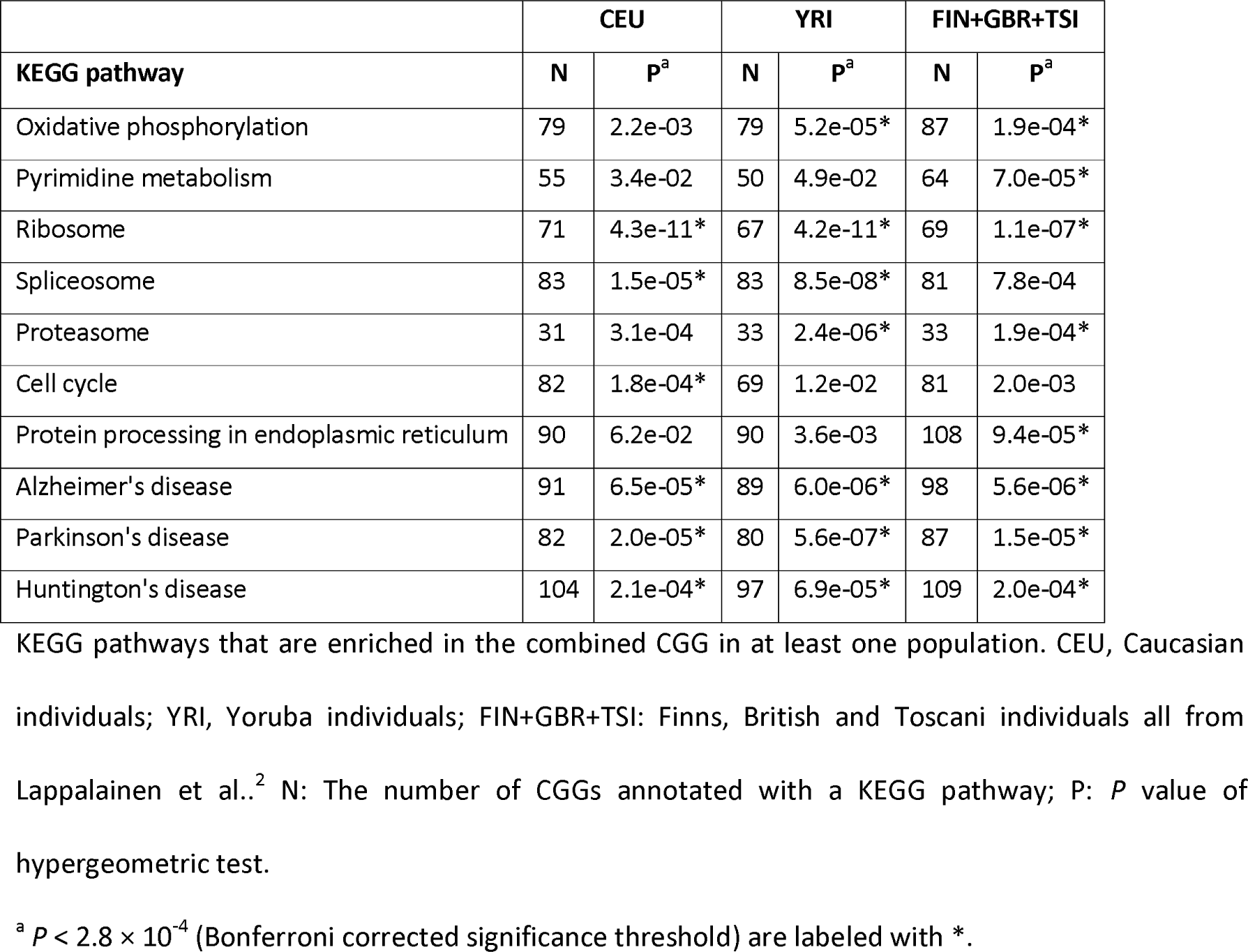
Enrichment of KEGG pathways in CGGs.

| KEGG pathway | CEU |  | YRI |  | FIN+GBR+TSI |  |
| --- | --- | --- | --- | --- | --- | --- |
|  | N | P<sup>a</sup> | N | P<sup>a</sup> | N | P<sup>a</sup> |
| Oxidative phosphorylation | 79 | 2.2e-03 | 79 | 5.2e-05* | 87 | 1.9e-04* |
| Pyrimidine metabolism | 55 | 3.4e-02 | 50 | 4.9e-02 | 64 | 7.0e-05* |
| Ribosome | 71 | 4.3e-11* | 67 | 4.2e-11* | 69 | 1.1e-07* |
| Spliceosome | 83 | 1.5e-05* | 83 | 8.5e-08* | 81 | 7.8e-04 |
| Proteasome | 31 | 3.1e-04 | 33 | 2.4e-06* | 33 | 1.9e-04* |
| Cell cycle | 82 | 1.8e-04* | 69 | 1.2e-02 | 81 | 2.0e-03 |
| Protein processing in endoplasmic reticulum | 90 | 6.2e-02 | 90 | 3.6e-03 | 108 | 9.4e-05* |
| Alzheimer's disease | 91 | 6.5e-05* | 89 | 6.0e-06* | 98 | 5.6e-06* |
| Parkinson's disease | 82 | 2.0e-05* | 80 | 5.6e-07* | 87 | 1.5e-05* |
| Huntington's disease | 104 | 2.1e-04* | 97 | 6.9e-05* | 109 | 2.0e-04* |

KEGG pathways that are enriched in the combined CGG in at least one population. CEU, Caucasian individuals; YRI, Yoruba individuals; FIN+GBR+TSI: Finns, British and Toscani individuals all from Lappalainen et al.<sup>2</sup> N: The number of CGGs annotated with a KEGG pathway; P: *P* value of hypergeometric test.
<sup>a</sup> $P < 2.8 \times 10^{-4}$ (Bonferroni corrected significance threshold) are labeled with \*.

### Expressions of spliceosome genes correlate with changes in splice patterns

Testing of phenotypic outcomes of variation in the CGGs enriched in particular pathways in most cases would require further case control or biochemical analyses, but the hypothesis that expression variation within the spliceosome set of genes results in changes to splice patterns can readily be tested by analysis of the alternative splicing in the same RNA-seq data that we have used in our analyses. Variation in mRNA levels will result in some variation in protein levels and Battle et al.^46^ showed the median Spearman’s correlation between LCL mRNA levels and protein levels across 62 Yoruban individuals for 4322 genes is 0.14. Even though this correlation is relatively weak the mean variation for the 83 spliceosome genes in the combined CGG is 3.2 fold for mRNA levels and 3.0 fold for protein levels (see Table S14). Given this variability we sought to ask if covariation in the mRNA levels of the spliceosome genes might contribute to changes in splicing. We focused upon the RNA-Seq data of 88 individuals from the Yoruban and Caucasian sample sets from Lappalainen et al.^2^ and we used the software MISO^35^ which uses annotations of alternative splicing events of human genes (see Materials and Methods). We calculated the PSI value for each alternative splicing event in the LCL. PSI values were calculated for two classes of alternative splicing: retained introns (Rl), where an intron is retained in the mRNA transcript relative to other transcripts from the same gene where the intron is removed, and skipped exons (SE) where an exon is removed by splicing in some but not all transcripts of the same gene. In the Yoruban samples MISO detects 1656 RI events in 1162 genes and 10100 SE events in 4824 genes, and for the 88 Caucasian samples there were 1507 RI events in 1100 genes and 9258 SE events in 4579 genes.

We then ask how variability of PSI values of RI and SE events might be attributed to the variability of the expressions of spliceosome genes. In total in the 88 Yoruban data set there are 83 (out of 107 with detectable mRNA levels) spliceosome genes in the combined CGG and for the Caucasian individuals there are 83 out of 110 detectable genes: 73 genes are shared in the combined CGG of both populations. We hypothesized that expressions of spliceosome genes may contribute to the variability of gene splicing, and also that variability of CGG from spliceosome genes should explain more variability of gene splicing than non-CGG due to their *trans* effects on gene splicing. To test this hypothesis, we regressed PSI values of individual splicing events on PC1 to PC20 eigenvectors of expressions of spliceosome genes combined. We observed that for Rl events using Yoruban and (Caucasian) RNA-seq data, CGG of spliceosome genes explain 29% (31%) of splicing variability on average, while non-CGG of spliceosome genes explain only 13% (17%) on average. This result suggests that 16% (14%) of splicing variability can be attributed to *trans* effects from expressions of CGG of spliceosome genes. Similarly, for SE events using Yoruban and (Caucasian) RNA-seq data, CGG of spliceosome genes explain 16% (17%) of splicing variability on average, while non-CGG of spliceosome genes explain only 7.6% (8.5%) on average, so that 8.4% (8.5%) of splicing variability can be attributed to *trans* effects from expressions of CGG of spliceosome genes. These data suggest that for both populations variation of CGG from the spliceosome pathway has a significant impact upon differential splicing of genes; the CGG’s expression variation accounts for an average of 16% of splicing variation for the RI events in the Yoruban data, and given the variation explained is the average R^2^ value for 1656 RI events this is a substantial influence.

## Discussion

In this paper, we have used multiple lines of evidence to show that correlation in mRNA levels has a biological rather than purely technical origin. In summary, we show this by using PCA of multiple sets of mRNA-Seq data to identify genes with correlated mRNA levels. We then use four lines of evidence to show covariation is unlikely to be due to batch effects: 1) they can be replicated within and between populations and across different analytic platforms including mRNA-Seq and microarray; 2) they contain an overabundance of cognate TFs bound to their census promoter regions and they are enriched for genes known to be sensitive to knockdown of single TFs; 3) the proteins they encode are more likely to physically interact with each other than expected; 4) they comprise groups of genes encoding shared functions more frequently than expected. These multiple lines of evidence lead us to believe that CGGs are indeed covarying under the influence of variation in their *trans*-acting controllers and that the cumulative *trans* effects can account for a significant proportion of individual genes’ expression variation.

## The properties of CGGs align with what is known about control of mRNA levels

The control of mRNA levels can occur at any point from initiation of transcription, to splicing, mRNA stability and degradation and these process all involve multiple protein and/or RNA interactions in *trans*.^47^ If we focus upon transcriptional control by TFs then any given gene can be controlled by multiple TFs and the action of any given TF can be activating or repressive conditional on proximity to promoter and/or interacting with or co-binding to other TFs. Variation of a TF can therefore be expected to influence sets of genes that based upon Cusanovich et al.^7^ can number from a few dozen to over 3000. However, the simultaneous variation of multiple TFs will necessarily have complicated outcomes conditional upon the overlap of influence; presently we have no understanding of the extent of these outcomes but it is likely that the mRNA of subsets of genes will be increased, decreased or remain unaltered in any one individual under the simultaneous variation of two TFs. Ultimately the combinatorial nature of the control of gene expression by TFs means that *trans* influences could influence anywhere from several thousand down to a single gene and that these effects could be significantly different both between individuals and between populations; this behaviour is indeed what we observe in our replication studies where we observe significant overlap, but not identity, of CGGs.

## How much variation is associated with combined *trans* influences?

Comparison of the variance sequentially removed by PCA for the two sets of genes reveals that CGG lose variance faster than non-CGG until PC9 when the reverse becomes true for the RNA-Seq data of Yoruban, the Caucasian and the British, Finns and Toscani populations from Lappalainen et al.^2^ The total variance of expressions of CGG accounted by the first 9 PCs is 63.8%, 62.2% and 64.3%, respectively and the total variance of expression of non-CGG accounted for by the first 9 PCs are 37.8%, 37.8% and 37.5%, respectively. Therefore, the difference in average variance lost between CGG and non-CGG is 26.0%, 24.4% and 26.8% for the 3 populations, respectively. It is reasonable that the difference reflects the effects of the shared variation which, by virtue of its scale, is recognized earlier in the PCA sequential analysis and suggests that at least 24% of mRNA level variation appears to be due to combined *trans* effects.

## Comparison to other studies

Goldinger et al.^16^ systematically studied the influence of PCA on mRNA levels and eQTLs using microarray data and tests of SNP association and heritability of expression. They concluded “*Most importantly, we show that the first 50 PCs, which have been removed in previously published articles to correct for batch effects* (*references omitted*), *contain*…… *a considerable proportion of genetic variation influencing gene expression*……”. We note that microarray and mRNA-Seq have quite different distributions of variance explained per PC: for example, PC1 explains a much greater proportion of variance from microarray data than it does from mRNA-Seq data. This means our findings cannot be directly related to those of Goldinger et al.^16^ both because the variance structures are very different and also because we can have no insight into the causation of shared influences— a necessary feature of our approach. However, our findings similarly suggest batch effects and biological signal can be confounded and techniques to de-noise mRNA-Seq data need to be applied very carefully. As we discuss above, the PEER software of Stegle et al.^27^ can effectively remove most of the covariation that underpins CGGs’ behavior.

The approach we have developed has advantages and disadvantages over present more gene centric eQTL approaches. The advantages are, firstly, the overall level of an mRNA is determined by many influences and it is the actual level that ultimately may be associated with biological outcome. In contrast, eQTL based approaches dissect the causative variations influencing mRNA levels and given these are individually small effects and subject to epistatic interactions this can be a challenging approach to detail the final mRNA level variation. Secondly, our approach does not generate the multiple testing problem of *trans* eQTL analyses and so can be applied to relatively small data sets. Thirdly, the correlated mRNA changes within functionally related genes may well prove to have a greater predictive effect upon phenotypic outcomes than the single *cis*- or *trans*-eQTL associations. The disadvantages to our approach are, firstly that we require mRNA level analysis of phenotypically relevant tissues (a problem which is increasingly shared by tissue specific eQTL analyses) and secondly that we do not gain mechanistic understanding at the level of association between DNA sequence variation and regulators or targets. We also recognize that the LCL resources we have used here are tissue culture cell lines and therefore subject to culture induced changes (see for example Yuan et al.^48^ and Choy et al.^49^) which may confound analysis.

## Spliceosome mRNA variation and alternative splicing

The finding that collective variation in the mRNA level of the spliceosome genes may account for 14–16% of the variation in alternative splicing of RI events enables us to ask if collective behavior of CGGs might have a stronger effect than variation in expression of individual genes. We compare the 14–16% collective influence to the influence of single spliceosome genes using a similar analysis framework. We calculated the average correlation between variation in the mRNA levels of individual genes and the splicing profile of individual RI events across the same individuals, and we observe correlations in both the Caucasian and the Yoruban data (see Table S15) that range from - 0. 24 to 0.19 and R^2^ of up to 0.065 with about 1/3 of the single gene correlations being positive; these figures replicate with a correlation between the two populations of 0.72. The very complex structure of CGGs covariation combined with the complex biochemistry of splicing events where coordinated changing levels of multiple proteins could modulate splicing activities, suggests that for any set of genes in a pathway their overall effect of expression variation is likely to enhance the impact of variation in individual genes. This highlights the interesting possibility that the control of gene expression in *trans* might have substantial overall effect on human phonotypes even if impact of individual genes is small.

## Alzheimer’s, Parkinson’s and Huntington’s disease and oxidative phosphorylation

The enrichment in the combined CGG for KEGG functions Alzheimer’s disease (62%, 62% and 68% of 158 KEGG annotated genes from Caucasian, Yoruban and British, Finns, Toscani populations respectively), Parkinson’s diseases (65%, 63% and 69% of 127 genes) and Huntington’s disease (59%, 55%, and 62% of 177 genes) is particularly striking (see Table S16 for details). In the Yoruban samples of Battle et al.^46^ the mRNA levels vary with a median of 3.0, 2.9 and 2.9 fold change and protein levels with a median of 4.2, 4.4 and 3.9 fold change for AD, PD and HD genes, respectively. The enrichment is driven in good measure by 58 genes functionally classified as oxidative phosphorylation which is itself an enriched function (62%, 62% and 68% of 128 genes). These 58 genes are detected as enriched in the combined CGG of all 3 populations and a further 12 oxidative phosphorylation genes are found in CGGs from one or two populations only. In total, an additional 17 genes involved in oxidative phosphorylation are found in one or more of the 3 disease functions in CGGs from one or more populations yielding a total of 88 genes involved in oxidative phosphorylation out of 265 genes in the combined list of genes involved in one or more disease categories. We also found 71 mitochondria localized genes that are shared with AD, PD and HD genes. The involvement of the mitochondrion in the etiology of all three diseases has been extensively documented (see Wang et al.,^50^ Franco-lborra et al.,^51^ Turner and Schapira^52^ for reviews of mitochondrial dysfunction and AD, PD and HD respectively and Biffi et al.^53^ for genetic evidence for involvement in AD) and our detection of correlated changes raises the question of whether this variation results in significant change to mitochondrial function and therefore an increased susceptibility to, or severity of, disease outcome. It has also been widely discussed (see Sun et al.^54^ for a recent review) that aging processes are associated with changes to mitochondrial function and properties. This suggests that identifying an effect of the natural covariation of genes involved in the oxidative phosphorylation pathway could be a significant contribution to understand the natural variation of human aging process.

For the remaining functional categories, pyrimidine metabolism, ribosome, spliceosome, proteasome, cell cycle, protein processing in endoplasmic reticulum, there is relatively little overlap with the exception of four proteins involved in ubiquitin conjugating and a ubiquitin ligase shared between protein processing in endoplasmic reticulum and Parkinson’s disease (see Ross et al.^55^ for a review of the role of ubiquitin and mitochondrial damage in PD and AD) and 10 RNA polymerase II subunits shared between pyrimidine metabolism and Huntington’s disease.

In each case testing of the relationship of variation in mRNA levels to ultimate biochemical phenotype will be challenging but we believe that we have defined some potential targets that could contribute to natural human variation that in turn might have significant impact upon our health.

## Acknowledgments

We would like to thank Teo Yik Ying for advice and insight. This work was supported by grants to P.F.R.L. from the National University of Singapore (NUS), through the Office of Deputy President (Research & Technology) and the Life Sciences Institute of NUS. Q.Y. was supported by a NUS Research Scholarship. Contributions to this work by R.B.H.W. were supported in part by the Australian National University.

## References

1. Nicolae, D.L., Gamazon, E., Zhang, W., Duan, S., Dolan, M.E., and Cox, N.J. (2010). Trait-associated SNPs are more likely to be eQTLs: annotation to enhance discovery from GWAS. PLoS Genet. 6, e1000888.

2. Lappalainen, T., Sammeth, M., Friedländer, M.R., Hoen, P.A., Monlong, J., Rivas, M.A., Gonzàlez-Porta, M., Kurbatova, N., Griebel, T., Ferreira, P.G., et al. (2013). Transcriptome and genome sequencing uncovers functional variation in humans. Nature 501, 506–511.

3. Albert, F.W., and Kruglyak, L. (2015). The role of regulatory variation in complex traits and disease. Nat. Rev. Genet. 16, 197–212.

4. Huan, T., Liu, C., Joehanes, R., Zhang, X., Chen, B.H., Johnson, A.D., Yao, C., Courchesne, P., O’Donnell, C.J., Munson, P.J., et al. (2015). A systematic heritability analysis of the human whole blood transcriptome. Hum. Genet. 134, 343–358.

5. Yu, H., Luscombe, N.M., Qian, J., and Gerstein, M. (2003). Genomic analysis of gene expression relationships in transcriptional regulatory networks. Trends Genet. 19, 422–427.

6. Lovén, J., Orlando, D.A., Sigova, A.A., Lin, C.Y., Rahl, P.B., Burge, C.B., Levens, D.L., Lee, T.I., and Young, R.A. (2012). Revisiting global gene expression analysis. Cell 151, 476–482.

7. Cusanovich, D.A., Pavlovic, B., Pritchard, J.K., and Gilad, Y. (2014). The functional consequences of variation in transcription factor binding. PLoS Genet. 10, e1004226.

8. Barrera, L.A., Vedenko, A., Kurland, J. V., Rogers, J.M., Gisselbrecht, S.S., Rossin, E.J., Woodard, J., Mariani, L., Kock, K.H., Inukai, S., et al. (2016). Survey of variation in human transcription factors reveals prevalent DNA binding changes. Science 351, 1450–1454.

9. Wu, C., Delano, D.L., Mitro, N., Su, S. V, Janes, J., McClurg, P., Batalov, S., Welch, G.L., Zhang, J., Orth, A.P., et al. (2008). Gene set enrichment in eQTL data identifies novel annotations and pathway regulators. PLoS Genet. 4, e1000070.

10. Rotival, M., Zeller, T., Wild, P.S., Maouche, S., Szymczak, S., Schillert, A., Castagné, R., Deiseroth, A., Proust, C., Brocheton, J., et al. (2011). Integrating genome-wide genetic variations and monocyte expression data reveals trans-regulated gene modules in humans. PLoS Genet. 7, e1002367.

11. Reiner, A.P., Hartiala, J., Zeller, T., Bis, J.C., Dupuis, J., Fornage, M., Baumert, J., Kleber, M.E., Wild, P.S., Baldus, S., et al. (2013). Genome-wide and gene-centric analyses of circulating myeloperoxidase levels in the charge and care consortia. Hum. Mol. Genet. 22, 3381–3393.

12. Brynedal, B., Choi, J.M., Raj, T., Bjornson, R., Stranger, B.E., Neale, B.M., Voight, B.F., and Cotsapas, C. (2017). Large-scale *trans*-eQTLs affect hundreds of transcripts and mediate patterns of transcriptional co-regulation. Am. J. Hum. Genet. 100, 581–591.

13. Cowley, M.J., Cotsapas, C.J., Williams, R.B.H., Chan, E.K.F., Pulvers, J.N., Liu, M.Y., Luo, O.J., Nott, D.J., and Little, P.F.R. (2009). Intra- and inter-individual genetic differences in gene expression. Mamm. Genome 20, 281–295.

14. Leek, J.T., Scharpf, R.B., Bravo, H.C., Simcha, D., Langmead, B., Johnson, W.E., Geman, D., Baggerly, K., and Irizarry, R. a (2010). Tackling the widespread and critical impact of batch effects in high-throughput data. Nat. Rev. Genet. 11, 733–739.

15. Chen, C., Grennan, K., Badner, J., Zhang, D., Gershon, E., Jin, L., and Liu, C. (2011). Removing batch effects in analysis of expression microarray data: an evaluation of six batch adjustment methods. PLoS One 6, el7238.

16. Goldinger, A., Henders, A.K., McRae, A.F., Martin, N.G., Gibson, G., Montgomery, G.W., Visscher, P.M., and Powell, J.E. (2013). Genetic and nongenetic variation revealed for the principal components of human gene expression. Genetics 195, 1117–1128.

17. Kolesnikov, N., Hastings, E., Keays, M., Melnichuk, O., Tang, Y.A., Williams, E., Dylag, M., Kurbatova, N., Brandizi, M., Burdett, T., et al. (2015). ArrayExpress update-simplifying data submissions. Nucleic Acids Res. 43, D1113–D1116.

18. Pickrell, J.K., Marioni, J.C., Pai, A. a, Degner, J.F., Engelhardt, B.E., Nkadori, E., Veyrieras, J.-B., Stephens, M., Gilad, Y., and Pritchard, J.K. (2010). Understanding mechanisms underlying human gene expression variation with RNA sequencing. Nature 464, 768–772.

19. Montgomery, S.B., Sammeth, M., Gutierrez-Arcelus, M., Lach, R.P., Ingle, C., Nisbett, J., Guigo, R., and Dermitzakis, E.T. (2010). Transcriptome genetics using second generation sequencing in a Caucasian population. Nature 464, 773–777.

20. Li, H., and Durbin, R. (2009). Fast and accurate short read alignment with Burrows-Wheeler transform. Bioinformatics 25, 1754–1760.

21. Li, H., Handsaker, B., Wysoker, A., Fennell, T., Ruan, J., Homer, N., Marth, G., Abecasis, G., and Durbin, R. (2009). The sequence alignment/map format and SAMtools. Bioinformatics 25, 2078–2079.

22. Morgan, M., Pagès, H., Obenchain, V., and Hayden, N. (2016). Rsamtools: Binary alignment (BAM), FASTA, variant call (BCF), and tabix file import. R package version 1.24.0.

23. Lawrence, M., Huber, W., Pagès, H., Aboyoun, P., Carlson, M., Gentleman, R., Morgan, M.T., and Carey, V.J. (2013). Software for computing and annotating genomic ranges. PLoS Comput. Biol. 9, e1003118.

24. Flicek, P., Ahmed, I., Amode, M.R., Barrell, D., Beal, K., Brent, S., Carvalho-Silva, D., Clapham, P., Coates, G., Fairley, S., et al. (2012). Ensembl 2013. Nucleic Acids Res. gks1236.

25. Stranger, B.E., Montgomery, S.B., Dimas, A.S., Parts, L., Stegle, O., Ingle, C.E., Sekowska, M., Smith, G.D., Evans, D., Gutierrez-Arcelus, M., et al. (2012). Patterns of cis regulatory variation in diverse human populations. PLoS Genet. 8, e1002639.

26. Csardi, G., and Nepusz, T. (2006). The igraph software package for complex network research. InterJournal, Complex Syst. 1695, 1–9.

27. Stegle, O., Parts, L., Piipari, M., Winn, J., and Durbin, R. (2012). Using probabilistic estimation of expression residuals (PEER) to obtain increased power and interpretability of gene expression analyses. Nat. Protoc. 7, 500–507.

28. Gerstein, M.B., Kundaje, A., Hariharan, M., Landt, S.G., Yan, K.-K., Cheng, C., Mu, X.J., Khurana, E., Rozowsky, J., Alexander, R., et al. (2012). Architecture of the human regulatory network derived from ENCODE data. Nature 489, 91–100.

29. Kanehisa, M., Goto, S., Furumichi, M., Tanabe, M., and Hirakawa, M. (2010). KEGG for representation and analysis of molecular networks involving diseases and drugs. Nucleic Acids Res. 38, D355–D360.

30. Carlson, M. (2013). org.Hs.eg.db: Genome wide annotation for Human (R package version 2.10.1).

31. Calvo, S.E., Clauser, K.R., and Mootha, V.K. (2015). MitoCarta2.0: an updated inventory of mammalian mitochondrial proteins. Nucleic Acids Res. 44, D1251-7.

32. Szklarczyk, D., Morris, J.H., Cook, H., Kuhn, M., Wyder, S., Simonovic, M., Santos, A., Doncheva, N.T., Roth, A., Bork, P., et al. (2017). The STRING database in 2017: quality-controlled protein-protein association networks, made broadly accessible. Nucleic Acids Res. 45, D362–D368.

33. Franceschini, A., Szklarczyk, D., Frankild, S., Kuhn, M., Simonovic, M., Roth, A., Lin, J., Minguez, P., Bork, P., and von Mering, C. (2013). STRING v9. 1: protein-protein interaction networks, with increased coverage and integration. Nucleic Acids Res. 41, D808–D815.

34. Kim, D., Pertea, G., Trapnell, C., Pimentel, H., Kelley, R., and Salzberg, S.L. (2013). TopHat2: accurate alignment of transcriptomes in the presence of insertions, deletions and gene fusions. Genome Biol. 14, R36.

35. Katz, Y., Wang, E.T., Airoldi, E.M., and Burge, C.B. (2010). Analysis and design of RNA sequencing experiments for identifying isoform regulation. Nat Methods 7, 1009–1015.

36. Qin, S., Kim, J., Arafat, D., and Gibson, G. (2013). Effect of normalization on statistical and biological interpretation of gene expression profiles. Front. Genet. 3, 160.

37. Fehrmann, R.S.N., Jansen, R.C., Veldink, J.H., Westra, H.-J., Arends, D., Bonder, M.J., Fu, J., Deelen, P., Groen, H.J.M., Smolonska, A., et al. (2011). *trans*-eQTLs reveal that independent genetic variants associated with a complex phenotype converge on intermediate genes, with a major role for the HLA. PLoS Genet. 7, e1002197.

38. Kanehisa, M., Goto, S., Sato, Y., Kawashima, M., Furumichi, M., and Tanabe, M. (2014). Data, information, knowledge and principle: back to metabolism in KEGG. Nucleic Acids Res. 42, D199–D205.

39. Allocco, D.J., Kohane, I.S., and Butte, A.J. (2004). Quantifying the relationship between coexpression, co-regulation and gene function. BMC Bioinformatics 5, 18.

40. Kasowski, M., Grubert, F., Heffelfinger, C., Hariharan, M., Asabere, A., Waszak, S.M., Habegger, L., Rozowsky, J., Shi, M., Urban, A.E., et al. (2010). Variation in transcription factor binding among humans. Science 328, 232–235.

41. Spitz, F., and Furlong, E.E.M. (2012). Transcription factors: from enhancer binding to developmental control. Nat. Rev. Genet. 13, 613–626.

42. Spivakov, M. (2014). Spurious transcription factor binding: Non-functional or genetically redundant? BioEssays 36, 798–806.

43. Farnham, P.J. (2009). Insights from genomic profiling of transcription factors. Nat. Rev. Genet. 10, 605–616.

44. Wang, J., Zhuang, J., Iyer, S., Lin, X., Whitfield, T.W., Greven, M.C., Pierce, B.G., Dong, X., Kundaje, A., Cheng, Y., et al. (2012). Sequence features and chromatin structure around the genomic regions bound by 119 human transcription factors. Genome Res. 22, 1798–1812.

45. Ideker, T., and Sharan, R. (2008). Protein networks in disease. Genome Res. 18, 644–652.

46. Battle, A., Khan, Z., Wang, S.H., Mitrano, A., Ford, M.J., Pritchard, J.K., and Gilad, Y. (2015). Impact of regulatory variation from RNA to protein. Science 347, 664–667.

47. Williams, R.B.H., Chan, E.K.F., Cowley, M.J., and Little, P.F.R. (2007). The influence of genetic variation on gene expression. Genome Res. 17, 1707–1716.

48. Yuan, Y., Tian, L., Lu, D., and Xu, S. (2015). Analysis of genome-wide RNA-sequencing data suggests age of the CEPH/Utah (CEU) lymphoblastoid cell lines systematically biases gene expression profiles. Sci. Rep. 5, 7960.

49. Choy, E., Yelensky, R., Bonakdar, S., Plenge, R.M., Saxena, R., De Jager, P.L., Shaw, S.Y., Wolfish, C.S., Slavik, J.M., Cotsapas, C., et al. (2008). Genetic analysis of human traits in vitro: drug response and gene expression in lymphoblastoid cell lines. PLoS Genet. 4, e1000287.

50. Wang, X., Wang, W., Li, L., Perry, G., Lee, H., and Zhu, X. (2014). Oxidative stress and mitochondrial dysfunction in Alzheimer–s disease. Biochim. Biophys. Acta - Mol. Basis Dis. 1842, 1240–1247.

51. Franco-Iborra, S., Vila, M., and Perier, C. (2016). The Parkinson Disease Mitochondrial Hypothesis: Where Are We at? Neurosci. 22, 266–277.

52. Turner, C., and Schapira, A.H. V (2010). Mitochondrial matters of the brain: The role in Huntington's disease. J. Bioenerg. Biomembr. 42, 193–198.

53. Biffi, A., Sabuncu, M.R., Desikan, R.S., Schmansky, N., Salat, D.H., Rosand, J., and Anderson, C.D. (2014). Genetic variation of oxidative phosphorylation genes in stroke and Alzheimer's disease. Neurobiol. Aging 35, 1956-e1.

54. Sun, N., Youle, R.J., and Finkel, T. (2016). The Mitochondrial Basis of Aging. Mol. Cell 61, 654–666.

55. Ross, J.M., Olson, L., and Coppotelli, G. (2015). Mitochondrial and ubiquitin proteasome system dysfunction in ageing and disease: Two sides of the same coin? Int. J. Mol. Sci. 16, 19458–19476.

